# Metabolic role of the LutACB operon in *Neisseria gonorrhoeae* resistance to neutrophil-mediated clearance

**DOI:** 10.1101/2025.07.10.664078

**Authors:** Ravisha Rawal, Jutta Schmidt, Werner Schmitz, Motaharehsadat Heydarian, Maximilian Klepsch, Thomas Rudel, Vera Kozjak-Pavlovic

**Affiliations:** Chair of Microbiology, University of Wuerzburg, Wuerzburg, Germany; Chair of Biochemistry and Molecular Biology, University of Wuerzburg, Wuerzburg, Germany; Perioperative, Acute, Critical Care and Emergency Medicine Section, Department of Medicine, University of Cambridge, UK

## Abstract

*Neisseria gonorrhoeae,* an obligate human pathogen, causes the sexually transmitted disease gonorrhoea and primarily colonizes the human genitourinary tract. An infection triggers a strong neutrophil-driven inflammatory response; however, viable gonococci are frequently recovered from neutrophil-rich patient exudates, indicating that these pathogens can survive neutrophil-mediated killing *in vivo*. To identify bacterial factors involved in this interaction, we screened a *N. gonorrhoeae* transposon library for mutants with reduced survival in polymorphonuclear leukocytes (PMNs). Among the identified candidates was NGFG_RS05085 (*lutB*), encoding an L-lactate oxidation FeS protein within the LutACB operon, which includes *lutA* (NGFG_RS05075), an FeS-binding protein, and *lutC*, the lactate utilisation protein C, (NGFG_RS05080). Deletion of *lutA*, *lutB*, or *lutC* did not significantly impair growth in standard culture medium or in minimal medium complemented with glucose, L-Lactate, or pyruvate. *lutB* and *lutC*, but not *lutA*, knockout significantly reduced bacterial survival within neutrophils compared to the wild-type strain. Polar effects of the mutations were ruled out by monitoring gene expression in deletion mutants. Metabolic profiling revealed disruption in nucleotide and glutathione synthesis, connecting the LutACB operon with sulfur metabolism, which could be required for a defence against neutrophil-derived reactive oxygen species (ROS). These findings highlight the important novel role of the LutACB operon, which connects lactate utilization and sulfur metabolism, and enables *N. gonorrhoeae* to withstand neutrophil-mediated oxidative stress during infection.

## Introduction

Gonorrhoea is a sexually transmitted disease caused by *Neisseria gonorrhoeae* (gonococcus or GC) with an estimated global incidence of nearly 87 million infections each year. Due to the emergence of extensive drug-resistance (Ohnishi *et al*, 2011; Quillin & Seifert, 2018), the World Health Organisation has designated Gc as a high-priority pathogen, and the Centers for Disease Control and Prevention (CDC) has classified it as an urgent antibiotic-resistant threat (Caméléna *et al*, 2024).

Whereas GC primarily infects the mucosal epithelium of the urogenital tract, it can also colonise other exposed mucosal surfaces, including rectum, pharynx, and conjunctivaDuring infection, gonococci must tackle a complex interplay of host immune factors, such as neutrophils, oxidative stress conditions, and nutrient availability, which influence bacterial survival and virulence (Hill *et al*, 2016; Seib *et al*, 2006; Walker *et al*, 2023). Despite this, GC persists at the infection site due to its adaptation to the human host and its ability to evade or manipulate host immune defences (Palmer & Criss, 2018; Quillin & Seifert, 2018).

Neutrophils, or polymorphonuclear leukocytes (PMNs), are the most abundant leukocytes in human blood and serve as the first line of defence against bacterial pathogens (Rosales, 2018; Witter *et al*, 2016). The colonization with GC triggers a strong inflammatory response (Quillin & Seifert, 2018) characterized by a significant influx of PMNs to the site of infection (Stevens & Criss, 2018). This neutrophil-driven response is a hallmark of gonococcal infections, playing a crucial role in both bacterial clearance and disease pathogenesis. PMNs use multiple antimicrobial strategies, including phagocytosis, oxidative burst (ROS production), degranulation, and the release of neutrophil extracellular traps (NETs) to neutralize and eliminate the invading bacteria (Palmer & Criss, 2018; Segal, 2005). Beyond direct microbial killing, neutrophils also play key roles in modulating inflammation through cytokine production and interactions with other immune cells (Château & Seifert, 2016).

For most bacterial infections, PMNs’ antimicrobial mechanisms effectively clear pathogens. However, GC has evolved strategies to evade PMN-mediated elimination, enabling persistent infection and contributing to the disease progression (Palmer & Criss, 2018). The bacterium employs antigenic and phase variation of surface pathogenicity factors, such as Opa proteins and pili, to escape antibody-mediated opsonization and recognition by phagocytes (Walker *et al*., 2023). Additionally, GC secretes nucleases such as Nuc to degrade neutrophil extracellular traps (NETs) (Juneau *et al*, 2015), and modulate phagolysosome trafficking within neutrophils, delaying the formation of mature degradative phagolysosomes (Johnson & Criss, 2013). During infection, GC is also exposed to reactive oxygen and nitrogen species generated by PMNs as part of the antimicrobial arsenal (Stohl & Seifert, 2006). Additionally, commensal *Lactobacillus* species in the vaginal flora produce hydrogen peroxide, which also contributes to the oxidative stress encountered by GC during colonization (Seib *et al*, 2005). To counteract these challenges, GC employs a manganese-based defence system to quench superoxide radicals and a glutathione-dependent system to neutralize nitric oxide (Kidd *et al*, 2005; Seib *et al*., 2005; Stohl *et al*, 2005; Tseng *et al*, 2001).

The metabolic adaptability of GC is a critical factor in its ability to withstand neutrophil-mediated killing during infection and contributes to its growing antimicrobial resistance (Ayala *et al*, 2024). The central carbon metabolism is tightly regulated, enabling bacteria to thrive even when the nutrients are limited or fluctuating. GC relies entirely on the human mucosal environment for nutrients, which closely links its metabolism to survival in the presence of host-derived stressors (Fu *et al*, 1989; Potter & Criss, 2024). Lactate, present in high amounts in the female genitourinary tract, but also a major metabolite released by neutrophils, can be efficiently used by GC as an energy source and can stimulate bacterial metabolism, further supporting bacterial ability to colonize the host and survive during neutrophil attack (Ayala & Shafer, 2019; Britigan *et al*, 1988; Exley *et al*, 2007). Mutants deficient in lactate dehydrogenases (LDHs) exhibit significantly reduced survival within neutrophils and impaired colonization of cervical epithelial cells, underscoring the importance of lactate utilization in gonococcal infections (Atack *et al*, 2014; Exley *et al*., 2007; Potter & Criss, 2024)

In this study, we performed a screen using a GC transposon mutant library (Remmele *et al*, 2014) to identify genes connected to the survival within neutrophils. Among these was lutB, a L-lactate oxidation FeS protein, which is a part of the previously characterised LutACB operon, traditionally associated with L-lactate utilisation (Chen *et al*, 2020). Our work, however, proposes an alternative function of this operon in GC survival upon PMN exposure. Upon deletion of the genes of the LutACB operon we observed impaired glutathione synthesis, which points to their function in sulphur metabolism and explains an increased susceptibility of deletion mutants to neutrophil antimicrobial defence. Our data suggest a so far unrecognised role of the LutACB operon in the metabolic adaptation of GC, enhancing its survival in the presence of neutrophils.

## Results

### A Tn5 transposon library screen identifies genes important for survival of GC in the presence of neutrophils

A genome-wide transposon mutant library of *N. gonorrhoeae* MS11 N2009 (Remmele *et al*., 2014) was used to perform a negative-selection screen in the PMN infection model. Three subsequent rounds of PMN infection yielded output libraries, which were analysed by high-throughput sequencing and compared to the respective input library to determine differentially represented transposon insertion sites, hence identifying GC genes important for survival in PMNs. The significantly underrepresented genes in the surviving bacteria were largely associated with the type IV pilus, which is critical for bacterial attachment and immune evasion; other hits included proteins linked to oxidative stress defence (e.g., catalase) and nucleic acid metabolism (e.g., ATP-dependent RNA helicase and MTA/SAH nucleosidase) (Supplementary table 1). Among the genes identified, NGFG_RS05085 (LutB) was of particular interest, as it encodes a LutB/LldF family L-lactate oxidation FeS protein that is potentially involved in lactate metabolism. Metabolising lactate, in turn, has been demonstrated to play a critical role in the survival of GC in neutrophils (Atack *et al*., 2014). LutB is part of an operon that also includes NGFG_RS05080 (LutC) and NGFG_RS05075 (LutA) (Figure 1A). This operon, previously characterized by Chen *et al*, is implicated in lactate utilization and represents an alternative L-lactate utilization pathway in *N. gonorrhoeae* in addition to the previously characterized L-lactate dehydrogenase LldD (Chen *et al*., 2020). Adjacent to the LutACB operon are NGFG_RS05070 (Multidrug efflux MFS Transporter, or MfSt) and NGFG_RS05090 (Type II toxin-antitoxin system toxin, or FitB), which flank the operon but are not directly involved in its regulatory structure (Figure 1A).

**Figure 1.**
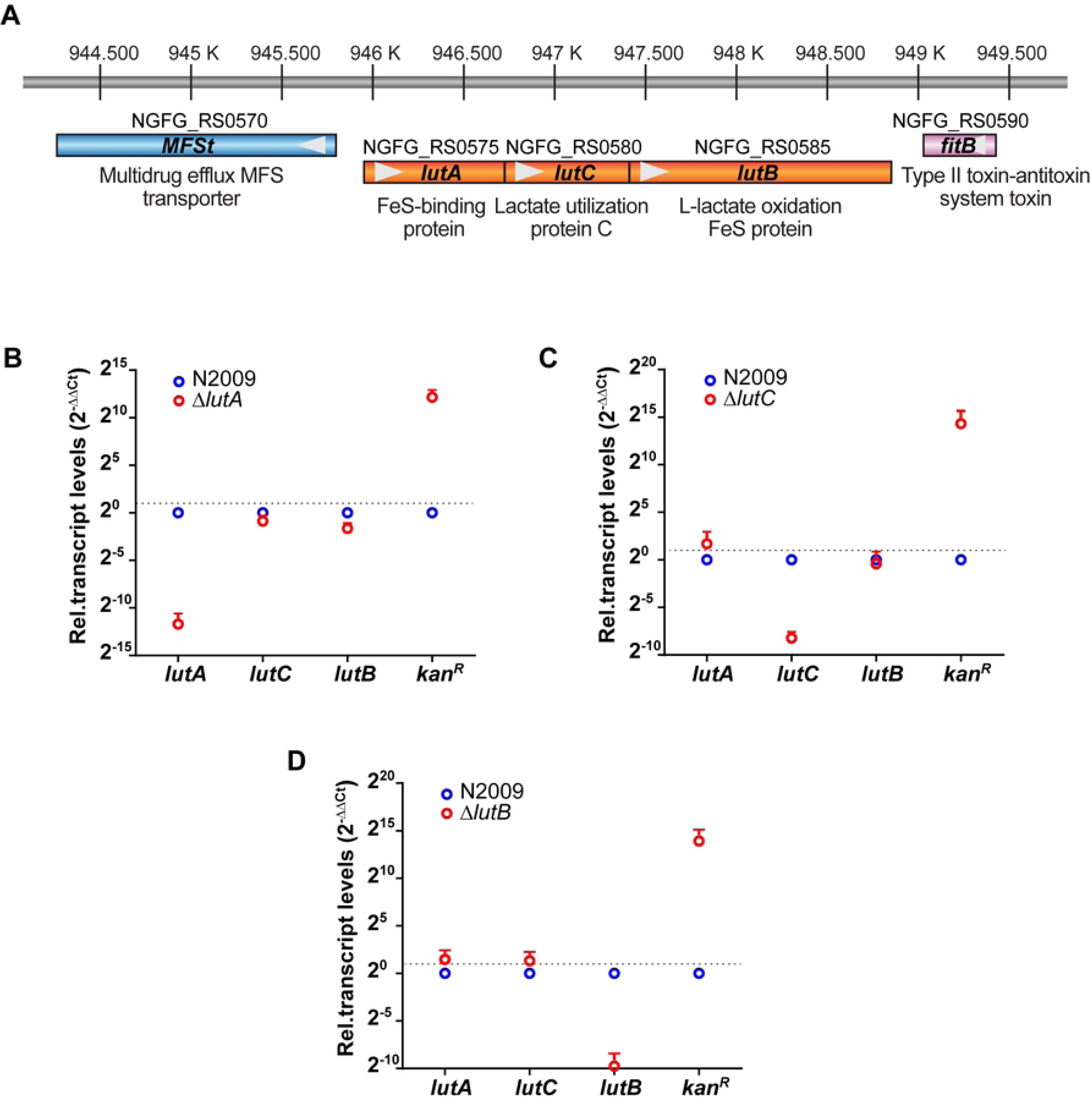
Deletion of genes of the LutACB operon does not affect expression of adjacent genes in *Neisseria gonorrhoeae*. (A) Schematic representation of the genomic organization of the LutACB operon and flanking genes. (B-D) Quantitative real-time PCR (RT-qPCR) analysis of relative gene expression levels in Δ*lutA*, Δ*lutB*, and Δ*lutC* compared to the wild-type strain MS11 (N2009). Transcript levels were normalized to 5S rRNA gene, and the fold change was calculated using the 2^−ΔΔCt^ method. The wild-type strain served as the reference. Each bar represents the mean ± SD from three independent experiments using cDNA synthesized from separate RNA preparations.

### Targeted gene knockouts of the LutACB operon and flanking genes show no polar effects

We constructed deletion mutants of the LutACB operon and flanking genes in *N. gonorrhoeae* MS11 (N2009) strain by replacing target genes with a kanamycin resistance cassette (Kan^R^). To assess potential polar effects of the gene disruption, the expression levels of relevant genes were measured by RT-qPCR, with normalization to the 5S rRNA gene. The knockout mutants demonstrated an increased Kan^R^ expression coupled with a reduction in the expression of the target gene mRNA, as expected (Figure 1B, C, D, Figure S1). Interestingly, the deletion of any member of the LutACB operon did not significantly affect expression of the remaining genes, indicating an absence of polar effects (Figure 1B, C, D). The deletion of the upstream and downstream genes for MFSt and FitB did not affect the expression of genes in the LutACB operon either (Figure S1).

To further investigate the functional role of these genes, we complemented the expression of *lutA, lutC,* and *lutB* by introducing the deleted gene under *pil*E promoter into the region between the lactate permease gene (NGFG_RS08030) and the aspartate aminotransferase gene (NGFG_RS08045). The RT-qPCR results confirmed that complementation successfully restored the expression of the deleted genes; however, the expression levels were somewhat higher than in the wild type (Figure S2).

### Deletion mutants show no growth defect in the rich medium or the chemically defined media supplemented with different carbon sources

We next analysed whether the gene knockout and complementation affect bacterial viability or replication. Bacteria were grown in rich PPM medium under standard conditions, and optical density (OD_550_) was measured at each hour over 5 hours. LutACB operon genes deletion and complementation mutants exhibited slower growth compared to the wild-type MS11 (N2009) bacteria, but the differences were not significant (Figure 2). In addition, there were no differences between the mutants and the wild type when growing bacteria in RPMI, a cell culture medium used in the neutrophil infection assays, although overall, bacteria grew more slowly than in the PPM medium (Figure S3).

**Figure 2.**
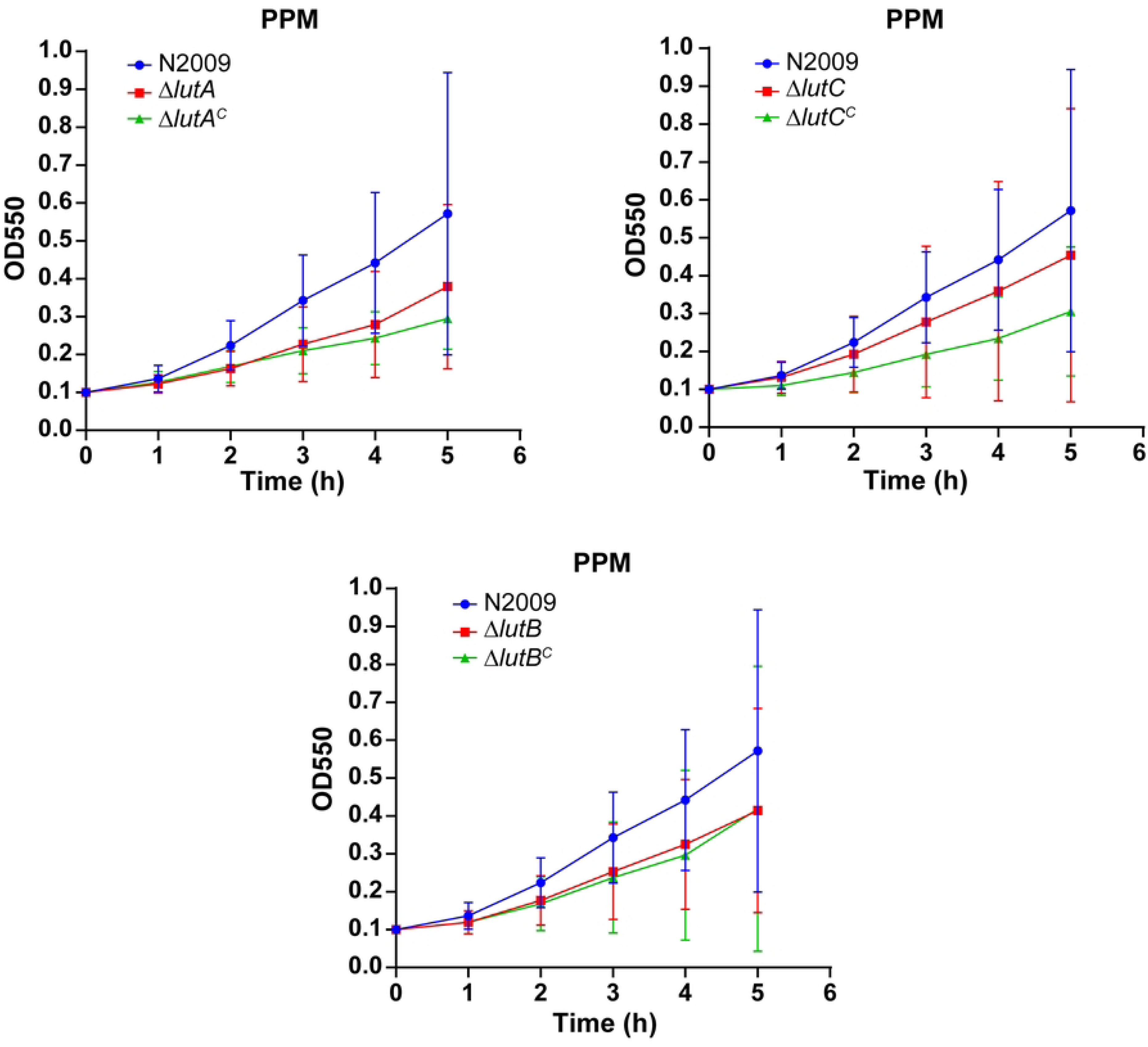
Deletion and complemented mutants exhibit no significant growth defect compared to wild type in PPM medium. Deletion mutants of *lutA, lutC*, and *lutB*, as well as the wild-type strain, were cultured in PPM supplemented with 1 % vitamin mix and 0.042 % NaHCO_3_. Mutants were grown under identical conditions, and optical density at 550 nm (OD550) was measured hourly over 5 h. The graphs show the mean value ± SD of three independent experiments.

We further conducted growth assays in chemically defined media supplemented with glucose, L-lactate, and pyruvate as carbon and energy sources. The knockout mutants of LutACB operon genes exhibited reduced growth rates compared to the wild type across all tested substrates; however, these differences were not statistically significant (Figure 3).

**Figure 3.**
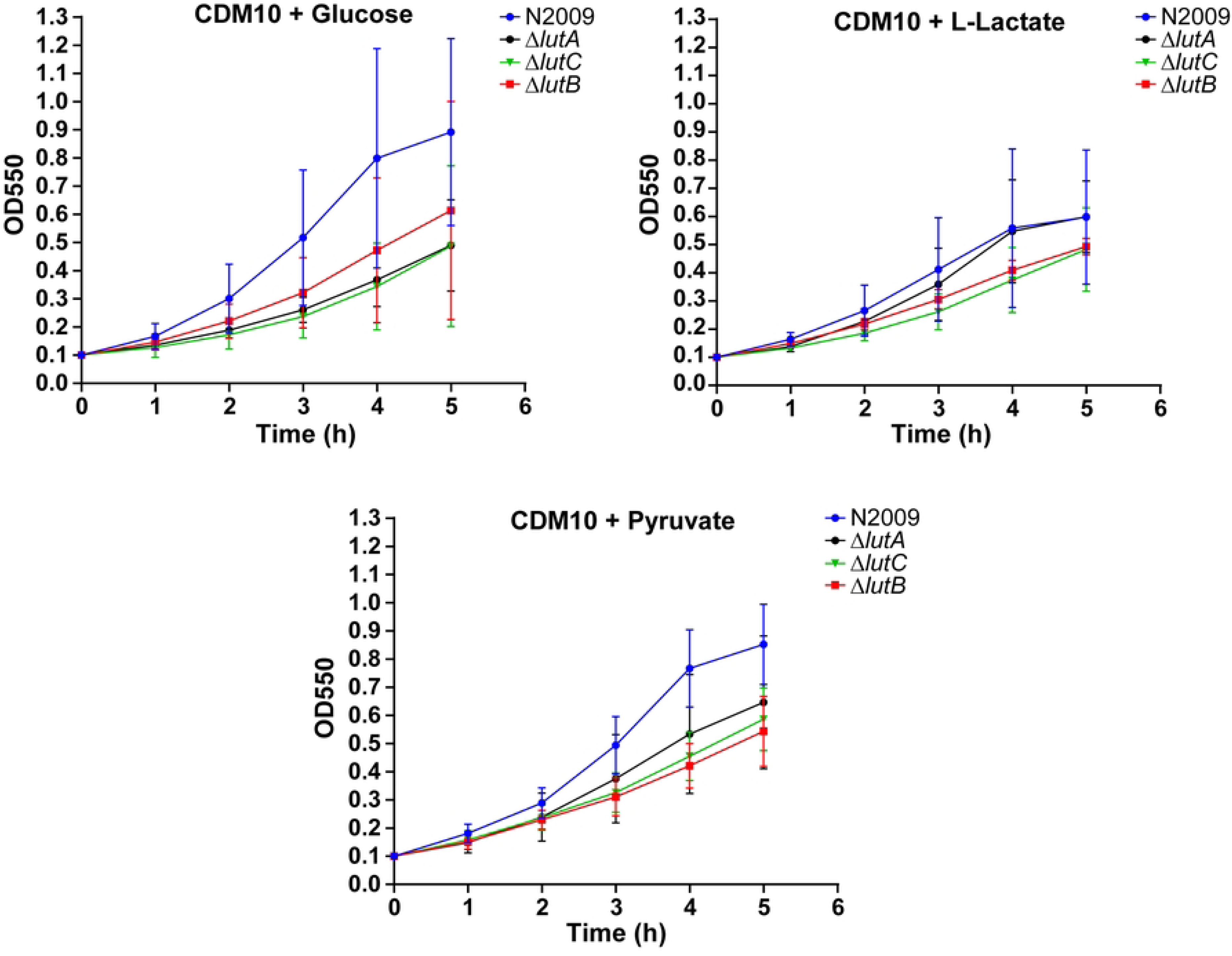
Growth curves of *N. gonorrhoeae* deletion strains in CDM supplemented with glucose, L-lactate, or pyruvate as the sole carbon source show no growth defect. Wild-type strain and deletion mutants of *lutA*, *lutC*, and *lutB* were cultured in chemically defined medium supplemented with either glucose, L-lactate, or pyruvate. The OD at 550 nm was measured hourly over a 5 h period to assess bacterial growth (OD550). The graphs represent mean ± standard deviation (SD) from three independent biological replicates.

When we tested the growth of Δ*fitB* and Δ*MFSt* bacteria, only Δ*MFSt* mutant growth was somewhat reduced in the PPM medium. In the RPMI medium, or in CDM supplemented with glucose, L-lactate, or pyruvate, the growth of both deletion mutants was comparable to that of the wild type (Figure S4, S5). For additional comparison, we generated deletion mutants of D-lactate dehydrogenase LdhD (NGFG_RS05005) and LctP (NGFG_RS07535), one of the two lactate permease genes. The growth of these deletion mutants was comparable to wild type in PPM medium, as well as the CDM medium supplemented with glucose or pyruvate, with slightly reduced growth in CDM + L-lactate. As expected, Δ*ldhD* GC failed to grow in CDM medium supplemented with D-lactate, whereas the growth of Δ*lctP* mutant was largely unaffected (Figure S6).

In conclusion, deletion of the members of the LutACB operon does not significantly affect the growth of GC in infection medium or in minimal medium with L-lactate as its only carbon and energy source.

### Deletion of LutB and LutC affects bacterial survival in the presence of neutrophils

To confirm the findings of the transposon library screen and assess whether the members of the LutACB operon are crucial for the survival of GC upon exposure to PMNs, we conducted a bacterial survival assay using primary PMNs isolated from whole blood at 1 h and 3 h post-infection (Figure 4). We could consistently isolate less viable GC from infected PMNs at the later time point, indicating bacterial killing by the neutrophils. The Δ*lutC* and Δ*lutB* mutants exhibited reduced survival compared to the wild-type strain at both time points, whereas for Δ*lutA*, this effect was not significant. Interestingly, only the complemented Δ*lutB^C^* mutant showed a significantly restored survival that approached wild-type levels. The complementation of LutC even increased the survival defect at 3 h post-infection (Figure 4). For the flanking genes, the Δ*fitB* mutant showed a modest reduction in survival at both time points; however, this reduction was not statistically significant. In contrast, the Δ*MFSt* deletion mutant did not show any significant difference in survival compared to the wild-type strain at either time point (Figure S7A). Δ*lctP* and Δ*ldhD* mutants showed some reduction in survival in the same assay, but due to the high variability of the measurements, this decrease was not significant (Figure S7B).

**Figure 4.**
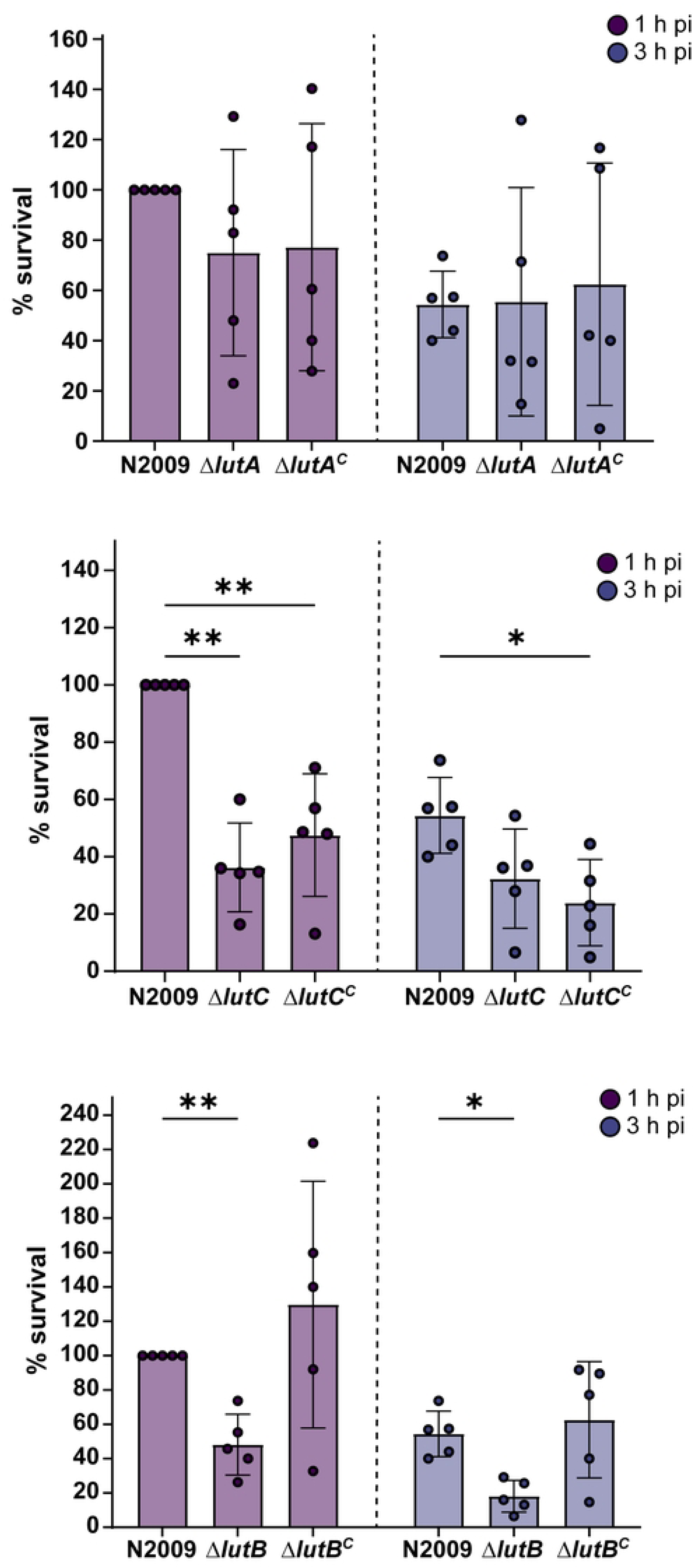
Deletion of *lutC* and *lutB* but not *lutA* affects bacterial survival upon exposure to neutrophils. Neutrophils isolated from donor blood were infected with Δl*utA*, Δ*lutC*, and Δ*lutB* deletion mutants and their respective complements. At 1 h and 3 h time points post-infection, infected neutrophils were lysed with saponin, and dilutions of lysate were plated on GC agar plates to assess the number of surviving bacteria. The data represent the mean ± SD of 5 independent experiments using different donors. The number of colonies was normalized to wild type at the 1 h time point. Statistical analysis was performed using two-way ANOVA. ^∗^p< 0.05, ^∗∗^p< 0.01

These findings indicate that *lutB* and *lutC* are important for the survival of GC upon neutrophil exposure. However, considering that the LutACB operon deletion mutants showed no significant defect when depending on L-lactate for growth (Figure 3), we wondered if this observed sensitivity to PMN killing can be attributed solely to a deficiency in L-lactate metabolism.

### Metabolomics and isotopologue profiling reveal altered redox homeostasis, sulphur metabolism and nucleotide biosynthesis in the knockout strain

To investigate metabolic alterations resulting from specific gene knockouts, we conducted a stable isotope labelling experiment using [U-¹³C₆] glucose, followed by isotopologue profiling. The results indicate that the bacteria mostly use Entner-Doudoroff pathway for pyruvate and acetyl-CoA generation, as well as diverting some labelled glucose into the pentose phosphate pathway for nucleotide synthesis, whereas the TCA cycle was largely inactive (Figure 5A, Supplementary Table 2). The only observed effect of LutB and LutC deletion was a decreased level of key purine metabolites, including adenosine and monophosphate (AMP and GMP) (Figure 5B, C).

**Figure 5.**
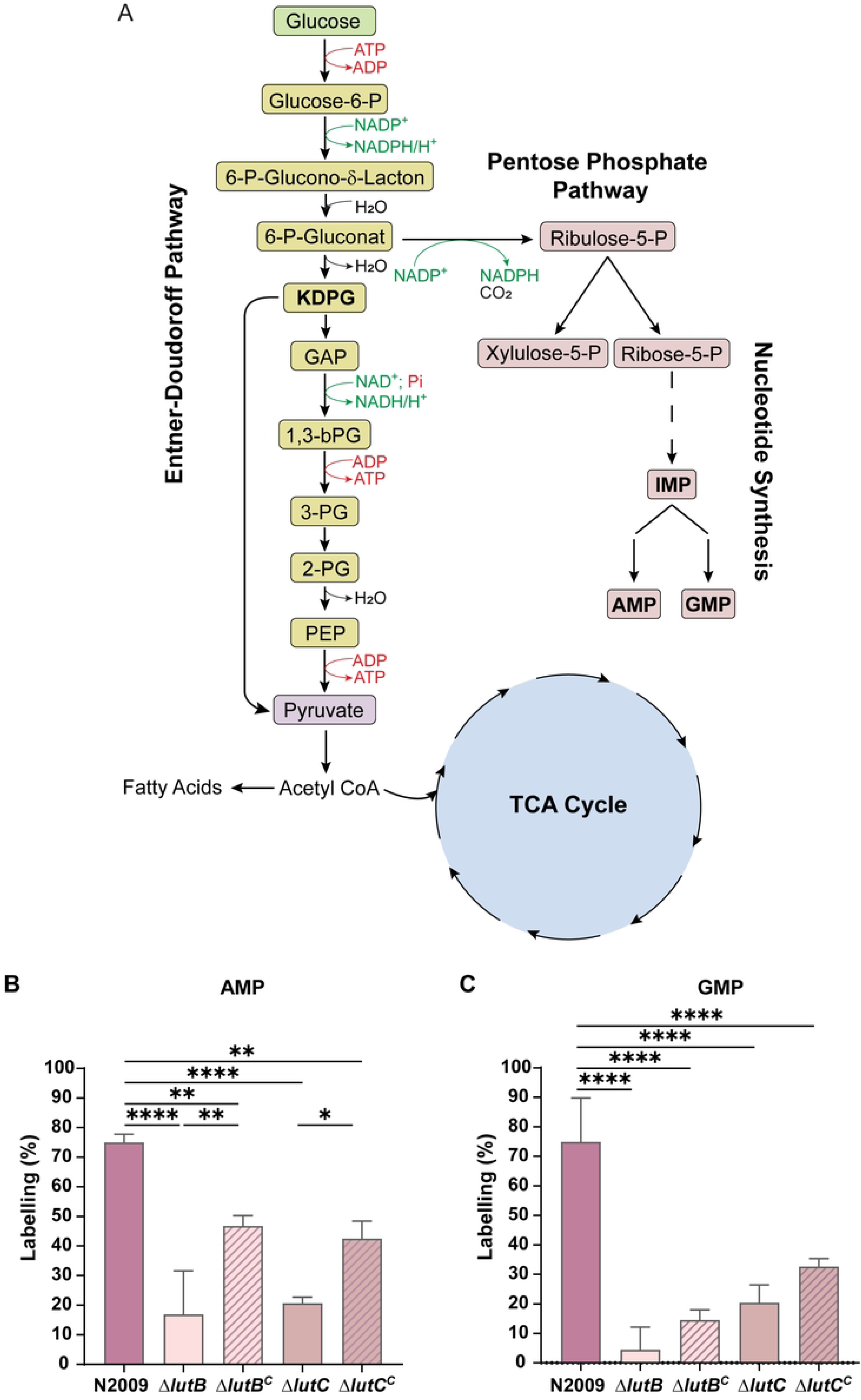
Disruption of *lutB* or *lutC* impairs AMP and GMP synthesis in *N. gonorrhoeae*. (A) Schematic representation of glucose metabolism in *N. gonorrhoeae*, with associated Entner-Doudoroff and Pentose Phosphate pathways. KDGP, 2-Keto-3-desoxy-6-phosphogluconat; GAP, Glycerinaldehyde-3-phosphate; 1,3-bPG, 1,3-Bisphosphoglycerate; 3-PG, 3-Phosphoglycerate; 2-PG, 2-Phosphoglycerate; PEP, Phosphoenolpyruvate; IMP, Inosine monophosphate; AMP, adenosine monophosphate; GMP, guanosine monophosphate; TCA, tricarboxylic acid. (B, C) Isotopologue profiling in wild type and deletion mutants of *lutB* and *lutC*, along with respective complemented mutants. The graphs show relative normalized peaks or labelling percentage of adenosine monophosphate (AMP) and guanosine monophosphate (GMP). The error bars represent the standard deviation calculated from three replicates. Statistical analysis was carried out by one-way ANOVA, followed by Tukey’s multiple comparison test, where asterisks indicate levels of significance (ns = not significant; *p<0.05; **p<0.01; ***p<0.001, ****p<0.0001).

Unlabelled total metabolite profiles of the wild-type, Δ*lutB*, Δ*lutC*, and their complements provided a broader perspective on the metabolic consequences of gene deletion. The comparison revealed alterations across multiple metabolic pathways (Figure 6, Supplementary Table 3). Considering that PMN survival defect for Δl*utB* could be complemented by *lutB* expression, we focused on those metabolites that were restored to wild-type levels in Δ*lutB^C^*. These metabolites were all a part of the sulfur metabolism (Figure S8) (Lauinger & Kaiser, 2021). Δl*utB* showed a slight decrease in the levels of S-adenosyl methionine (SAM), which were restored to even higher levels upon complementation (Figure 7A). Most pronounced, however, was the accumulation of cystine and cysteine (Figure 7B, C), as well as the reduction in glutathione (GSH) and its oxidised form, glutathione disulfide (GSSG), in both Δl*utB* and Δ*lutC*. This could be in part restored by complementation of Δl*utB*, but complementation of Δ*lutC* exacerbated the effect (Figure 7D, E). In conclusion, deletion of two members of the LutACB operon leads to an increase in cystine and cysteine levels, with a simultaneous decrease in GSH levels, pointing to a reduction in GSH synthesis or its increased degradation. Since GSH is one of the main compounds protecting cells from oxidative damage, the reduction in its levels upon *lutB* and *lutC* deletion could explain the increased sensitivity of the mutant GC to killing by neutrophils.

**Figure 6.**
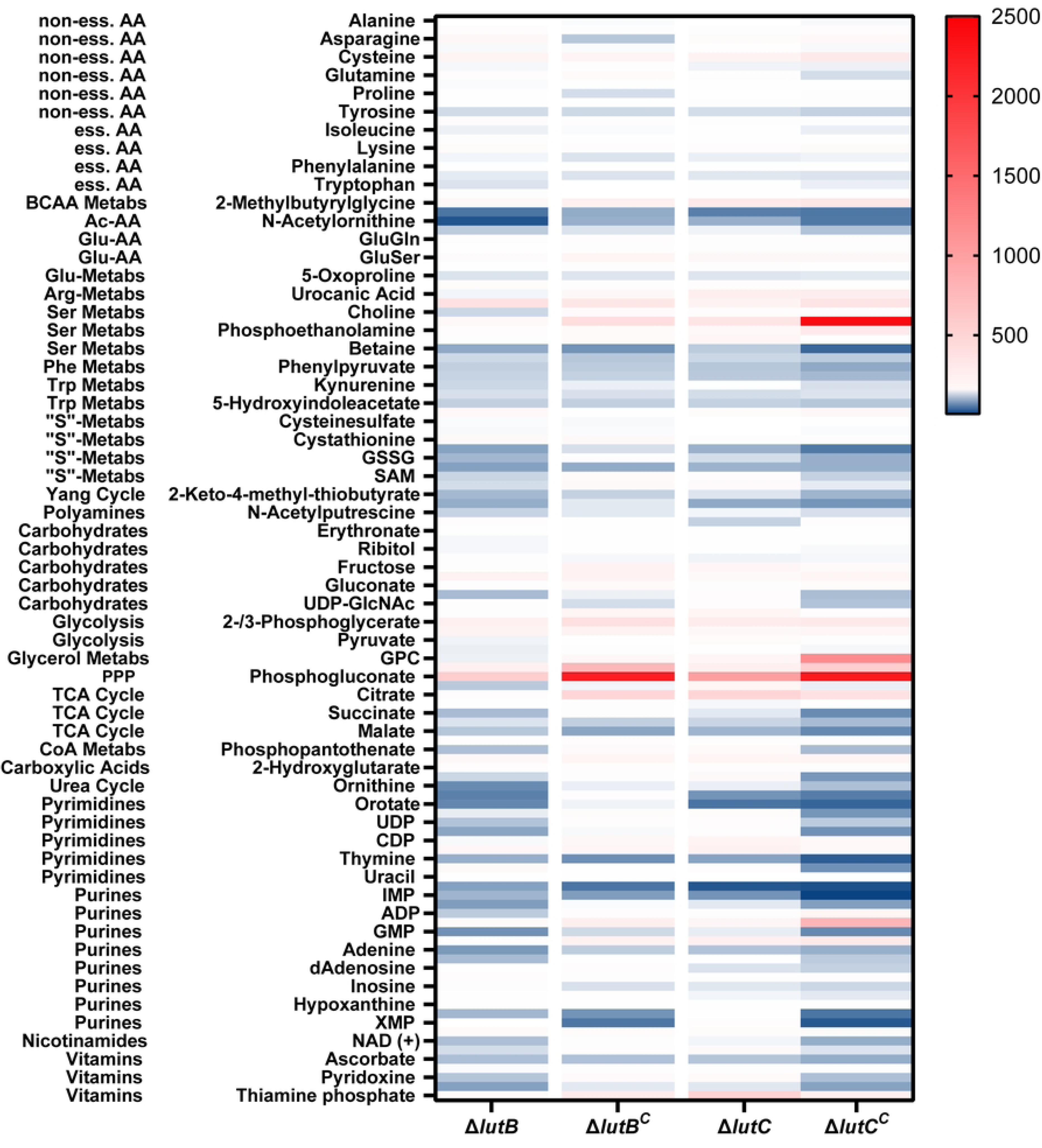
Heatmap of metabolite abundance in Δ*lutB*, Δ*lutB^C^*, Δ*lutC*, and Δ*lutC^C^* strains cultured in CDM supplemented with unlabelled glucose. The heatmap shows the relative concentrations of metabolites normalized to the wild type. Colour intensity indicates fold change in metabolite abundance. Supporting data is available in Supplementary Table 3.

**Figure 7.**
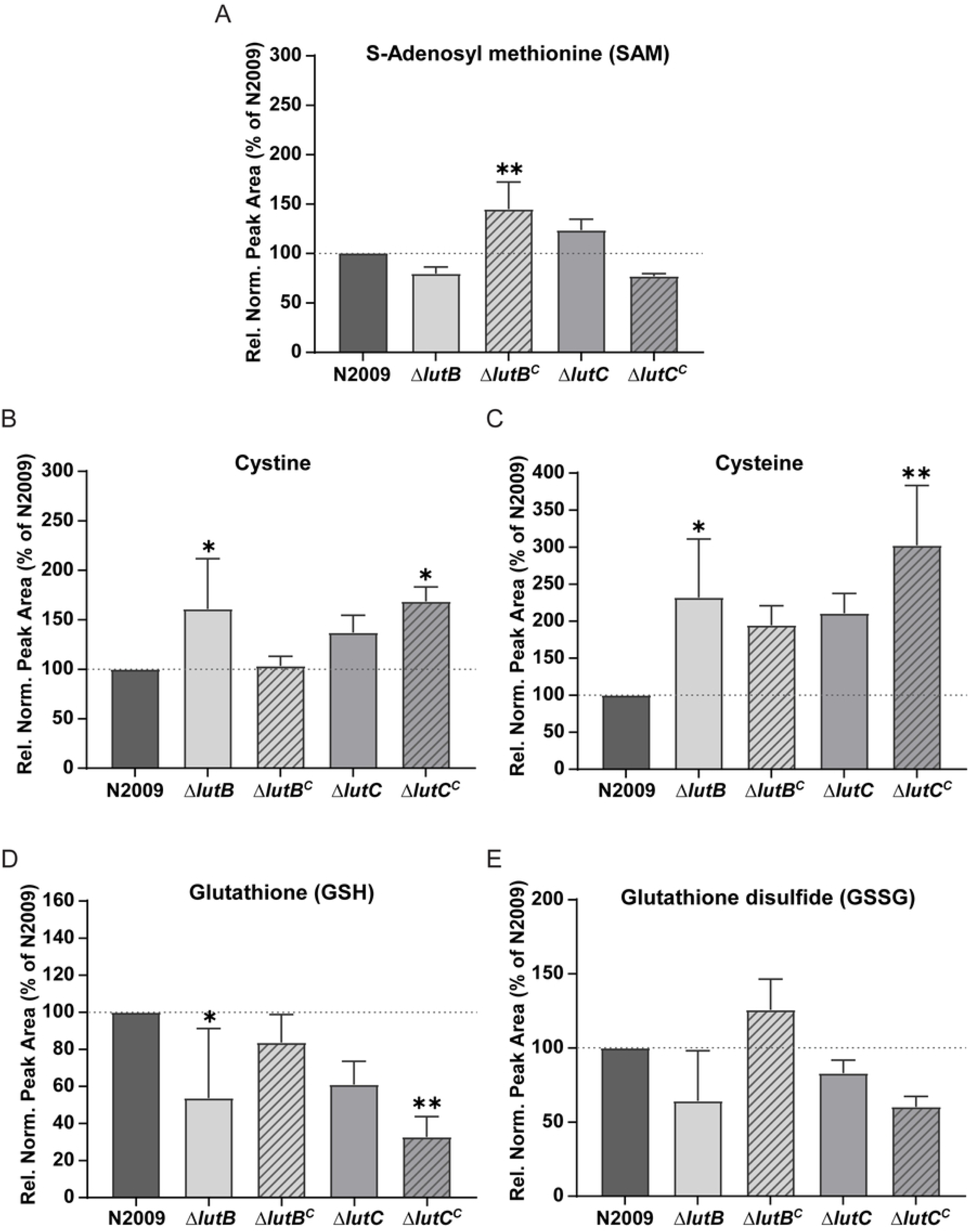
Quantitative analysis of sulfur-containing metabolites in Δ*lutB* and Δ*lutC* deletion and complemented mutants show accumulation of cystine and cysteine and reduction in SAM, GSH, and GSSG. The graphs represent normalized peak areas relative to wild-type values (percentage) for S-adenosyl methionine (SAM), cysteine, cystine, glutathione (GSH), and glutathione disulfide (GSSG). Metabolites were measured for Δ*lutB*, Δ*lutB^C^*, Δ*lutC*, and Δ*lutC^C^* mutants cultured in CDM supplemented with unlabelled glucose. Data are the mean ± standard deviation of three biological replicates. Statistical analysis was done using one-way ANOVA with Dunnett’s multiple comparisons test (*p<0.05; **p<0.01).

## Discussion

PMNs are a central component of the innate immune system and constitute the primary cellular defence against GC. Despite the array of potent oxidative and non-oxidative antimicrobial mechanisms deployed by PMNs, including the production of ROS, proteases, and antimicrobial peptides, GC demonstrates a remarkable ability to survive within this hostile environment (Criss *et al*, 2009; Seib *et al*., 2006).

Our study demonstrates that deletion of genes within the LutACB operon significantly reduces GC survival in the presence of neutrophils, while only mildly impairing the growth in defined media containing various carbon sources. The absence of strong growth defects in the Δ*lutA*, Δ*lutB*, and Δ*lutC* mutants when cultured in CDM supplemented with glucose, L-lactate, or pyruvate suggests that the presence of redundant L-lactate utilization pathways in *N. gonorrhoeae* (Chen *et al*., 2020) is sufficient to support bacterial growth. This agrees with a previous report, which indicated that LutACB did not play a role in lactate oxidation (Ibranovic, 2015) and implies that the LutACB operon in GC has a role different than its homologs in other bacteria (Chai *et al*, 2009), serving an important function during immune challenge, particularly in the oxidizing environment within neutrophils. The operon may contribute to metabolic adaptation, redox balance, or resistance to host-derived stresses, underscoring a potential role in virulence and intracellular persistence rather than general growth.

During infection, GC encounters significant oxidative stress from multiple sources. The bacterium is exposed to reactive oxygen and nitrogen species as a consequence of localised inflammatory responses in the mucosal surfaces it infects (Seib *et al*., 2006). A primary source of this oxidative stress comes from neutrophils, which generate ROS (such as superoxide, hydrogen peroxide, and hydroxyl radicals) as part of their antimicrobial defence mechanisms. Additionally, the activity of commensal lactobacilli in the vaginal flora contributes to the oxidative stress (Potter & Criss, 2024).

Our metabolic study of LutACB operon deletion mutants revealed alterations in nucleotide biosynthesis and levels of sulfur metabolites, supporting the role of this operon in cysteine metabolism and redox homeostasis. Isotope tracing with [U-¹³C₆] glucose demonstrated reduced ^13^C incorporation into AMP and GMP in the Δ*lutC* and Δl*utB*, indicating altered *de novo* nucleotide synthesis. These changes likely reflect broader metabolic stress and compromised energy homeostasis. The observed reduction in SAM (Figure 7A) could also, in part, explain the purine synthesis defect, considering its role in this process (Figure S8). Such a metabolic bottleneck may compromise the bacterium’s ability to resist neutrophil-mediated killing, as efficient nucleotide synthesis is essential for DNA repair and regulation of stress responses (LeCuyer *et al*, 2010; Stohl & Seifert, 2006).

Importantly, unlabelled total metabolite analysis revealed alterations in levels of multiple metabolites. Of these, the changes in sulfur-containing metabolites could be complemented by re-expression of LutB and corresponded to the neutrophil survival phenotype, highlighting a potentially critical role for sulfur metabolism in the physiological adaptation of GC to PMN-induced stress.

Glutathione (GSH), a tripeptide thiol, serves as an important cellular antioxidant, playing a crucial role in maintaining the redox homeostasis and bacterial defence against oxidative stress (Hicks & Mullholland, 2018). The reduced form (GSH) and the oxidized form (GSSG) together maintain the cellular redox balance. Cysteine, a precursor for GSH synthesis, is of particular importance to GC, which relies on the thiosulfate and cystine uptake for cysteine production. This dependence makes cysteine acquisition and metabolism vital for the pathogen’s survival (Hicks & Mullholland, 2018). In our study, both Δ*lutB* and Δl*utC* mutants showed elevated levels of cysteine and cystine, but reduced levels of GSH and GSSG compared to the wild-type MS11 (N2009). This metabolic shift suggests either a breakdown in the pathway converting cysteine to GSH or increased GSH degradation, which could influence the susceptibility to neutrophil-mediated killing (Johnson & Criss, 2011; Reniere, 2018). The complementation of *lutB* partially restored GSH amounts, indicating its importance for the maintenance of GSH levels. How this is achieved, we still do not fully understand, but the presence of the FeS cluster in LutB supports its role in redox reactions that could be important for GSH synthesis. Further studies should focus on the connection between the metabolism of thiosulfate, methionine, and cysteine and LutACB operon. Interestingly, the Δ*lutC^C^* complementation mutant showed even higher levels of cysteine and cystine, as well as lower GSH and GSSG levels than Δ*lutC*, which corresponded to the increased killing of Δ*lutC^C^* mutant by neutrophils at 3 h post-infection (Figure 4). The complementation was achieved by introducing *lutC* under a strong *pilE* promotor, which led to much higher expression of LutC in the Δ*lutC^C^* and lowered the expression of LutB (Figure S2B). This could explain the observed negative effect of the overproduction of LutC, indicating that a fine balance between the individual proteins encoded by the LutACB operon is required for the proper functioning of the metabolic processes in which they play a role.

The changes in SAM levels provided an additional insight into the metabolic alterations associated with the LutACB operon. As a key metabolite linking methylation reactions and sulphur metabolism, SAM indirectly influences cellular redox status by modulating cysteine and GSH synthesis, in addition to being connected to purine biosynthesis (Liu *et al*, 2022; Ouyang *et al*, 2020). In the Δ*lutB* mutant, the reduced SAM levels indicate impaired synthesis or increased consumption, likely due to disruption in the methionine biosynthesis pathway or its conversion to SAM (Berney *et al*, 2015). Complementation of *lutB* not only restored but overcompensated SAM biosynthesis, possibly due to a regulatory feedback mechanism. In contrast, Δ*lutC* showed moderately increased SAM levels, while its complementation led to a reduction in SAM amounts. These patterns underscore the complexity of interconnected metabolic networks, where compensatory mechanisms can inadvertently exacerbate metabolic imbalances. The increased level of SAM in the Δ*lutC* mutant may reflect impaired SAM utilization or altered feedback regulation, resulting in reduced consumption, accumulation of SAM, and a downstream increase in cysteine synthesis (Zhang *et al*, 2022). However, a block in GSH synthesis may prevent effective use of this excess cysteine. *lutC* is located centrally within the LutACB operon, and, as already discussed, the complementation using the strong *pilE* promoter may have disrupted the coordinated expression of operon genes. This could have caused metabolic imbalance due to altered protein stoichiometry, potentially interfering with pathway function (Wang *et al*, 2021), which highlights the importance of coordinated operon expression for maintaining metabolic homeostasis (Del Duca *et al*, 2023).

Together, our findings reveal that, contrary to its proposed role in lactate oxidation, the LutACB operon plays a role in sulfur metabolism and maintenance of the redox balance, which are crucial for GC survival within neutrophils and resistance to innate immune clearance.

## Materials and Methods

### Bacterial strains and growth conditions

The *N. gonorrhoeae* mutants used in this study were derived from the wild-type strain N2009 (PorB_IA_, Pili^+^, Opa^-^, cat<porBIA< >ermC) (Faulstich *et al*, 2013), the derivative strain of MS11 (GenBank accession number NC_022240.1). Bacteria were cultivated for 14 to 16 h at 37 °C in a humidified 5% CO_2_ atmosphere on GC agar plates (Oxoid, Wesel, Germany) supplemented with 1 % vitamin mix (Bauer *et al*, 2017). PPM containing 1 % vitamin mix and 0.04 % (w/v) NaHCO_3_ was used for liquid cultures (Per 1 l of dH_2_O: proteose peptone no. 3 [15 g], soluble starch [1 g], 11 % (w/w) K_2_HPO4 [4 g], 3 % (w/w) KH_2_PO4 [1 g], 14 % (w/w) NaCl [5 g]). Media were supplemented with kanamycin at final concentrations of 40 µg/ml when necessary.

### Transposon library neutrophil survival screen

Tn5 mutant library of *N. gonorrhoeae* MS11 (N2009) was generated as described before (Remmele *et al*., 2014). For the neutrophil survival screen, 10 μl of the library glycerol stock was diluted in 1 ml of PPM medium and cultured on 10 GC agar plates (20 cm diameter) at 37 °C with 5 % CO₂ for 16–18 h. Colonies were pooled in serum free RPMI to generate the INPUT library, which was used for infection and stored in part as glycerol stocks. 1.2 × 10⁷ of neutrophils isolated from donor blood were seeded on two 6-well plates (1 × 10⁶ cells/well), then centrifuged at 1000 g for 5 min to support neutrophil attachment, and infected at an MOI 30. Plates were centrifuged at 600 g for 3 min to synchronise infection and incubated at 37 °C for the total of 2 h. After 1 h, wells were washed three times with RPMI, followed by another hour of incubation. Surviving bacteria were recovered by lysing cells with 1% saponin, scraping, and plating the lysates on 24 GC agar plates (20 cm diameter). After 18–20 h incubation at 37 °C/5% CO₂, colonies were pooled in RPMI. The recovered bacteria were either stored or used for subsequent infection rounds. In total, three consecutive infections, yealding OUTPUT1, 2, and 3, were performed in triplicate. The resulting INPUT and OUTPUT3 libraries were sequenced with an Illumina HiSeq 3000 (150 bp reads, 20 million reads per sample) and analysed as already described (Remmele *et al*., 2014). Primers used are listed in Supplementary Table 4.

### DNA manipulations and genetic techniques

Upstream and downstream homologous fragments flanking the gene of interest (GOI) were amplified from N2009 genomic DNA and were fused to the kanamycin resistance cassette using overlap PCR. The resulting construct was transformed into piliated MS11 (N2009) to create the knockout mutants via homologous recombination (Kellogg *et al*, 1968). Each mutant strain was confirmed by PCR using forward and reverse gene- and Kan^R^ cassette–specific primers. Primers used are described in Supplementary Table 4.

For complementation, relevant genes fused with the *pilE* promotor were synthesized by a commercial provider (GeneArt, Invitrogen, Regensburg, Germany) and supplied within the pMA-RQ plasmid. The genes were recloned into a pPilE-mCherry-Spec (a kind gift from the group of Berenike Maier) plasmid using FseI and PacI restriction sites to exchange the mCherry reporter. Upon transformation of the resulting plasmid into *N. gonorrhoeae* recipient deletion mutant, the gene of interest was inserted between the *N. gonorrhoeae* lactate permease gene (NGFG_RS08030) and aspartate aminotransferase gene (NGFG_RS08045) by homologous recombination. Successful integration was confirmed by PCR and sequencing.

### Growth assay in defined medium

Growth in chemically defined medium was carried out in CDM10 (Dyer et al. 1987) with minor modifications (glucose, 2 g/l; L-lactate 2 g/l, sodium pyruvate, 2 g/l). Glucose was replaced by [U-^13^C_6_] glucose (a kind gift from gift group of Eisenreich, TU Munich), (2 g/l; 11.1 mM) for metabolic labelling experiments. The culture of *N. gonorrhoeae* was diluted to OD_550_ 0.1, and the bacteria were grown under aerobic conditions (37 °C, 120 rpm). OD_550_ was measured hourly for a period of 5 h.

### Quantitative gene expression studies

*N. gonorrhoeae* wild type, deletion, and complemented strains were grown to an OD_550_ of 0.5 in PPM. RNA was extracted in the mid-exponential growth phase using miRNeasy Micro kit (Qiagen, Hilden, Germany), followed by DNase 1 treatment to remove genomic DNA contamination (TURBO DNA-free, ThermoFisher Scientific, Hessen, Germany). Complementary DNA (cDNA) was synthesized from 1 µg of DNase-treated, RNase-free RNA using RevertAid First Strand cDNA Synthesis Kit (ThermoFisher, Hessen, Germany), using random hexamer primers. Reverse transcription quantitative PCR (RT-qPCR) was carried out using SYBR Green-based detection on StepOne Plus real-time PCR systems (Life Technologies, ThermoFisher Scientific, Hessen, Germany). Each 20 µl reaction mixture included diluted cDNA, GreenMasterMix (2X) High ROX (Gennaxon, Ulm, Germany), and 18 pM of each primer. Amplification conditions consisted of an initial denaturation at 95 °C for 10 min, followed by 40 cycles at 95 °C for 15 s and 60 °C for 60 s, ending with a dissociation curve analysis. At least three experiments were performed in triplicate (if not stated otherwise) with independently prepared cDNA, and relative gene expression levels were normalised to 5S rRNA gene using the 2^−ΔΔCT^ method (Livak & Schmittgen, 2001) in Excel (Microsoft). The primers used are listed in Supplementary Table 4. Raw data is available in Supplementary Table 5.

### Neutrophil killing assay

Killing assays were performed using human neutrophils. PMNs were isolated from healthy donors as already described (Moldovan *et al*, 2025). Bacterial suspensions were made in RPMI and grown (37 °C, 120 rpm shaking) to mid-logarithmic phase (OD₅₅₀ ≈ 0.4). All strains were then diluted in RPMI to equalise OD_550_ values, ensuring equal bacterial input for neutrophil infection. To promote adhesion, 2 × 10⁵ neutrophils were added to 12-well plates, centrifuged (5 min, 1,000 rpm), and infected with the respective strain at MOI 50. Until infection, plates were incubated at 37 °C with 5% CO₂. Cells were lysed with 1 % saponin (Sigma, Darmstadt, Germany), and serial dilutions of lysates were plated on GC agar, followed by incubation (37 °C, 5% CO₂) to count viable colony-forming units. The survival percentage was determined by normalising to the number of colonies of the wild-type strain at 1 h post-infection.

### Sample preparation for isotopologue profiling

Precultures were prepared for *N. gonorrhoeae* wild-type, deletion, and complemented strains in PPM medium and grown up to an OD_550_ of 0.4. The samples were washed and diluted to the same OD_550_ using carbon source-free CDM10 medium. Cultures were incubated at 37 °C with shaking at 120 rpm for 15 minutes. Subsequently, either unlabelled glucose or [U-^13^C_6_] glucose was added, followed by continued incubation under the same conditions. Samples supplemented with [^13^C_6_] glucose were harvested after 5 minutes, while those with unlabelled glucose were harvested after 60 min. The pellets were flash frozen in liquid nitrogen and resuspended in 500 μl of cold methanol/water (80:20, v/v). This was followed by amino acid and metabolite extraction and subsequent LC-MS analysis.

### Isotopologue profiling

Water-soluble metabolites were extracted with 500 μl ice-cold MeOH/H_2_O (80/20 v/v) containing 0.01 μM lamivudine and 10 μM each of D4-succinate, D5-glycine, D2-glucose, and 15N-glutamate as std. compounds (Sigma-Aldrich, St. Louis, USA). After centrifugation of the resulting homogenates, supernatants were evaporated in a rotary evaporator (Savant, Thermo Fisher Scientific, Waltham, USA). Dry sample extracts were redissolved in 150 μl of 5 μM NH_4_OAc in CH_3_CN/H_2_O (50/50, v/v). 15 μl of supernatant was transferred to LC vials. For LC-MS analysis, 3 μl of each sample was applied to an XBridge Premier BEH Amide (2.5 μm particles, 100× 2.1 mm) UPLC-column (Waters, Dublin, Ireland). Metabolites were separated with Solvent A, consisting of 5 mM NH_4_OAc in CH_3_CN/H_2_O (40/60, v/v) and solvent B, consisting of 5 mM NH_4_OAc in CH_3_CN/H_2_O (95/5, v/v) at a flow rate of 200 μl/min at 45 °C by LC using a DIONEX Ultimate 3000 UHPLC system (Thermo Fisher Scientific, Bremen, Germany). A linear gradient starting after 2 min with 100 % solvent B decreasing to 0 % solvent B within 23 min, followed by 17 min 0 % solvent B and a linear increase to 100 % solvent B in 1 min. Recalibration of the column was achieved by 7 min prerun with 0 % solvent B before each injection. Ultrapure H_2_O was obtained from a Millipore water purification system (Milli-Q Merck Millipore, Darmstadt, Germany). HPLC-MS solvents, LC-MS NH_4_OAc, standards, and reference compounds were purchased from Merck.

The resulting residue was resuspended in 20 μl pyridine buffer (86 ml H_2_O, 10.8 ml 6M HCl, 8.6 ml pyridine) containing 0.5 M each of BHA and ECD. After 1 h at RT, derivatized metabolites were extracted with 300 μl acetoacetate. The extract was washed with 300 μl H_2_O, evaporated under a stream of nitrogen at 45 °C and redissolved in 200 μl CH_3_CN/H_2_O (50/50, v/v) containing 0.1 % FA.

For LC-MS analysis, 3 μl of each sample was applied to an Atlantis Premier BEH C18 AX (2.5 μm particles, 100×2.1 mm) UPLC-column (Waters, Dublin, Ireland). Metabolites were separated with Solvent A, consisting of 0.1 % FA in CH_3_CN/H_2_O (5/95, v/v) and solvent B consisting of 0.1 % FA in CH_3_CN/H_2_O (90/10, v/v) at a flow rate of 200 μl/min at 45 °C by LC using the above-mentioned UHPLC system. A linear gradient starting after 2 min with 10 % solvent B increasing to 100 % solvent B within 20 min was followed by 10 min 100 % solvent B and a linear decrease to 10 % solvent B in 2 min. Recalibration of the column was achieved by 5 min prerun with 10 % solvent B before each injection.

All MS analyses were performed on a high-resolution Q Exactive mass spectrometer equipped with a HESI probe (Thermo Fisher Scientific, Bremen, Germany) in alternating positive- and negative full MS mode with a scan range of 69.0 1000 m/z at 70 K resolution and the following ESI source parameters: sheath gas: 30, auxiliary gas: 1, sweep gas: 0, aux gas heater temperature: 120 °C, spray voltage: 3 kV, capillary temperature: 320 °C, S-lens RF level: 50. XIC generation and signal quantitation was performed using TraceFinder V 5.1 (Thermo Fisher Scientific, Bremen, Germany) integrating peaks which corresponded to the calculated monoisotopic metabolite masses (MIM +/- H+ ± 3 mMU).

## Statistical analysis

Data were analysed using GraphPad Prism Software (Version 10.2.3). For statistical analysis, at least three biological replicates were performed (unless sample size is indicated for each graph). All data are presented as means with standard deviation (SD). P-values ≤0.05 were considered significant. Analysis of variance (ANOVA) was performed to determine whether the group means were significantly different from each other. For the time-dependent bacterial survival assay, a mixed model ANOVA with Dunnett’s multiple comparison test was performed to deduce individual differences. Details are provided in the figure legend of the corresponding figure.

## Acknowledgments

We would like to thank L. Speth, E. Maier, and K. Pekárková for technical assistance and B. Maier for the kind gift of plasmid. This work was supported by the German Research Foundation GRK2157 to TR and VK-P, as well as by the State of Bavaria SCIENTIA career support program to RR.

## Ethics statement

Venous blood was collected from healthy adult donors following informed consent. The use of human blood and neutrophils in this study was approved by the Ethics Committee of the University of Würzburg (approval number 300/21).

## Supplementary figure legends

**Figure S1. Deletion of *MFSt* and *fitB* does not affect LutACB operon gene expression.** Relative expression levels of *MFSt* and *fitB* genes, as well as of the LutACB operon genes, were measured using RT-qPCR in their respective knockout strains compared to wild type. Gene expression was normalized to 5S rRNA gene and calculated using 2^−ΔΔCt^ method. Data represents mean ± SD of three independent experiments.

**Figure S2. RT-qPCR analysis of gene expression in deletion and complemented strains.** The relative expression of the target gene in deletion mutants of *lutA, lutC*, and *lutB*, along with their complementation mutants, was measured and compared to the wild-type strain. Gene expression was normalized to the 5S rRNA housekeeping gene, and relative transcript levels were calculated using the 2^−ΔΔCt^ method. Data represent values of a single experiment.

**Figure S3. Growth curves in RPMI medium for the deletion and complemented mutants of the LutACB operon show no growth defect compared to wild type.** *lutA, lutC* and *lutB* deletion and complemented mutants as well as the wild type were cultured in RPMI under identical conditions, and optical density at 550 nm (OD550) was measured hourly over a 5 h period.

**Figure S4. Deletion of *MFSt* or *fitB* does not affect bacterial growth.** Optical density at 550 nm (OD550) was measured for Δ*MFSt* and Δ*fitB* mutants in PPM or RPMI medium every hour over 5 h. Graphs represent mean ± SD of three independent experiments.

**Figure S5. Δ*MFSt* and Δf*itB* grow comparable to wild type in CDM medium supplemented with either glucose, L-lactate, or pyruvate as the sole carbon source.** Wild type bacteria or Δ*MFSt* and Δ*fitB* mutants were cultured in chemically defined CDM medium supplemented with either glucose, L-lactate, or pyruvate. The OD at 550 nm (OD550) was measured each hour over a 5 h period to assess bacterial growth. The experiment was conducted in triplicate, and the graph represents mean ± standard deviation (SD).

**Figure S6. Deletion of ldhD impairs bacterial growth in CDM medium supplemented with D-lactate.** The growth was assessed for wild-type strain, as well as the deletion mutants of lactate permease *lctP* and D-lactate dehydrogenase *ldhD* in rich PPM medium, as well as in the defined CDM medium supplemented with either glucose, L-lactate, pyruvate, or D-lactate. Bacterial growth was monitored by measuring OD at 550 nm (OD550) every hour over a period of 5 h. Each data point represents the mean ± standard deviation (SD) from three independent experiments.

**Figure S7. Survival of *MFSt, fitB, lctP* and *ldhD* deletion mutants upon exposure to human neutrophils is not significantly affected.** Freshly isolated primary neutrophils were infected with wild-type strain MS11 (N2009), as well as Δ*MFSt*, Δ*fitB*, Δ*lctP* and Δ*ldhD* mutants, and bacterial survival was assessed at 1 h and 3 h post-infection. CFU/ml were calculated and normalized to the numbers for wild type at 1 h post-infection to determine percentage of surviving bacteria. Data represent the mean ± standard deviation (SD) from three independent experiments using neutrophils from different donors.

**Figure S8. Schematic representation of key points in sulfur metabolism.** The scheme shows the methionine cycle, in which S-adenosyl methionine (SAM) and S-adenosyl-L-homocysteine (SAH) are synthesized from methionine. Homocysteine serves as a precursor for cysteine synthesis, which, together with glutamate and glycine, serves to produce glutathione (GSH), and glutathione disulfide (GSSG). GSH is degraded through γ-glutamyl cycle to yield cysteine, among others.

## References

Atack JM, Ibranovic I, Ong C-LY, Djoko KY, Chen NH, vanden Hoven R, Jennings MP, Edwards JL, McEwan AG (2014) A Role for Lactate Dehydrogenases in the Survival of *Neisseria gonorrhoeae* in Human Polymorphonuclear Leukocytes and Cervical Epithelial Cells. J Infect Dis 210: 1311–1318

Ayala JC, Balthazar JT, Shafer WM (2024) Transcriptional responses of *Neisseria gonorrhoeae* to glucose and lactate: implications for resistance to oxidative damage and biofilm formation. mBio 15: e0176124

Ayala JC, Shafer WM (2019) Transcriptional regulation of a gonococcal gene encoding a virulence factor (L-lactate permease). PLOS Path 15: e1008233

Bauer S, Helmreich J, Zachary M, Kaethner M, Heinrichs E, Rudel T, Beier D (2017) The sibling sRNAs NgncR_162 and NgncR_163 of *Neisseria gonorrhoeae* participate in the expression control of metabolic, transport and regulatory proteins. Microbiology 163: 1720–1734

Berney M, Berney-Meyer L, Wong K-W, Chen B, Chen M, Kim J, Wang J, Harris D, Parkhill J, Chan J et al (2015) Essential roles of methionine and S-adenosylmethionine in the autarkic lifestyle of *Mycobacterium tuberculosis*. PNAS 112: 10008–10013

Britigan BE, Klapper D, Svendsen T, Cohen MS (1988) Phagocyte-derived lactate stimulates oxygen consumption by Neisseria gonorrhoeae. An unrecognized aspect of the oxygen metabolism of phagocytosis. J Clin Invest 81: 318–324

Caméléna F, Mérimèche M, Brousseau J, Mainardis M, Verger P, Le Risbé C, Brottet E, Thabuis A, Bébéar C, Molina J-M et al (2024) Emergence of Extensively Drug-Resistant *Neisseria gonorrhoeae*, France, 2023. Emerg Infect Dis 30: 1903

Chai Y, Kolter R, Losick R (2009) A Widely Conserved Gene Cluster Required for Lactate Utilization in *Bacillus subtilis* and Its Involvement in Biofilm Formation. J Bacteriol 191: 2423–2430

Château A, Seifert HS (2016) *Neisseria gonorrhoeae* survives within and modulates apoptosis and inflammatory cytokine production of human macrophages. Cell Microbiol 18: 546–560

Chen NH, Ong CY, O’Sullivan J, Ibranovic I, Davey K, Edwards JL, McEwan AG (2020) Two Distinct L-Lactate Dehydrogenases Play a Role in the Survival of *Neisseria gonorrhoeae* in Cervical Epithelial Cells. J Infect Dis 221: 449–453

Criss AK, Katz BZ, Seifert HS (2009) Resistance of *Neisseria gonorrhoeae* to non-oxidative killing by adherent human polymorphonuclear leucocytes. Cell Microbiol 11: 1074–1087

Del Duca S, Semenzato G, Esposito A, Liò P, Fani R (2023) The Operon as a Conundrum of Gene Dynamics and Biochemical Constraints: What We Have Learned from Histidine Biosynthesis. Genes (Basel) 14

Exley RM, Wu H, Shaw J, Schneider MC, Smith H, Jerse AE, Tang CM (2007) Lactate acquisition promotes successful colonization of the murine genital tract by *Neisseria gonorrhoeae*. Infect Immun 75: 1318–1324

Faulstich M, Böttcher J-P, Meyer TF, Fraunholz M, Rudel T (2013) Pilus Phase Variation Switches Gonococcal Adherence to Invasion by Caveolin-1-Dependent Host Cell Signaling. PLOS Path 9: e1003373

Fu HS, Hassett DJ, Cohen MS (1989) Oxidant stress in *Neisseria gonorrhoeae*: adaptation and effects on L-(+)-lactate dehydrogenase activity. Infect Immun 57: 2173–2178

Hicks JL, Mullholland CV (2018) Cysteine biosynthesis in *Neisseria* species. Microbiology 164: 1471–1480

Hill SA, Masters TL, Wachter J (2016) Gonorrhea - an evolving disease of the new millennium. Microb Cell 3: 371–389

Ibranovic I, 2015. Lactate dehydrogenases from *Neisseria gonorrhoeae*: molecular analysis and role in host cell/bacterial interactions, School of Chemistry and Molecular Biosciences. The University of Queensland, Queensland, Australia.

Johnson MB, Criss AK (2011) Resistance of *Neisseria gonorrhoeae* to neutrophils. Front Microbiol 2: 77

Johnson MB, Criss AK (2013) Neisseria gonorrhoeae phagosomes delay fusion with primary granules to enhance bacterial survival inside human neutrophils. Cell Microbiol 15: 1323–1340

Juneau RA, Stevens JS, Apicella MA, Criss AK (2015) A thermonuclease of Neisseria gonorrhoeae enhances bacterial escape from killing by neutrophil extracellular traps. J Infect Dis 212: 316–324

Kellogg DS, Cohen IR, Norins LC, Schroeter AL, Reising G (1968) *Neisseria gonorrhoeae* II. Colonial Variation and Pathogenicity During 35 Months In Vitro. J Bacteriol 96: 596–605

Kidd SP, Potter AJ, Apicella MA, Jennings MP, McEwan AG (2005) NmlR of *Neisseria gonorrhoea*e: a novel redox responsive transcription factor from the MerR family. Mol Microbiol 57: 1676–1689

Lauinger L, Kaiser P (2021) Sensing and Signaling of Methionine Metabolism. Metabolites 11: 83

LeCuyer BE, Criss AK, Seifert HS (2010) Genetic characterization of the nucleotide excision repair system of *Neisseria gonorrhoeae*. J Bacteriol 192: 665–673

Liu Q, Lin B, Tao Y (2022) Improved methylation in *E. coli* via an efficient methyl supply system driven by betaine. Metab Eng 72: 46–55

Livak KJ, Schmittgen TD (2001) Analysis of Relative Gene Expression Data Using Real-Time Quantitative PCR and the 2−ΔΔCT Method. Methods 25: 402–408

Moldovan A, Wagner F, Schumacher F, Wigger D, Kessie DK, Rühling M, Stelzner K, Tschertok R, Kersting L, Fink J et al (2025) *Chlamydia trachomatis* exploits sphingolipid metabolic pathways during infection of phagocytes. mBio: e0398124

Ohnishi M, Golparian D, Shimuta K, Saika T, Hoshina S, Iwasaku K, Nakayama S, Kitawaki J, Unemo M (2011) Is *Neisseria gonorrhoeae* initiating a future era of untreatable gonorrhea?: detailed characterization of the first strain with high-level resistance to ceftriaxone. Antimicrob Agents Chemother 55: 3538–3545

Ouyang Y, Wu Q, Li J, Sun S, Sun S (2020) S-adenosylmethionine: A metabolite critical to the regulation of autophagy. Cell Prolif 53: e12891

Palmer A, Criss AK (2018) Gonococcal Defenses against Antimicrobial Activities of Neutrophils. Trends Microbiol 26: 1022–1034

Potter AD, Criss AK (2024) Dinner date: *Neisseria gonorrhoeae* central carbon metabolism and pathogenesis. Emerg Top Life Sci 8: 15–28

Quillin SJ, Seifert HS (2018) *Neisseria gonorrhoeae* host adaptation and pathogenesis. Nat Rev Microbiol 16: 226–240

Remmele CW, Xian Y, Albrecht M, Faulstich M, Fraunholz M, Heinrichs E, Dittrich MT, Müller T, Reinhardt R, Rudel T (2014) Transcriptional landscape and essential genes of *Neisseria gonorrhoeae*. Nucleic Acids Res 42: 10579–10595

Reniere ML (2018) Reduce, Induce, Thrive: Bacterial Redox Sensing during Pathogenesis. J Bacteriol 200: 10.1128/jb.00128-00118

Rosales C (2018) Neutrophil: A Cell with Many Roles in Inflammation or Several Cell Types? Front Physiol 9: 113

Segal AW (2005) How neutrophils kill microbes. Ann Rev Immunol 23: 197–223

Seib KL, Simons MP, Wu H-J, McEwan AG, Nauseef WM, Apicella MA, Jennings MP (2005) Investigation of Oxidative Stress Defenses of *Neisseria gonorrhoeae* by Using a Human Polymorphonuclear Leukocyte Survival Assay. Infec Immun 73: 5269–5272

Seib KL, Wu HJ, Kidd SP, Apicella MA, Jennings MP, McEwan AG (2006) Defenses against oxidative stress in *Neisseria gonorrhoeae*: a system tailored for a challenging environment. Microbiol Mol Biol Rev 70: 344–361

Stevens JS, Criss AK (2018) Pathogenesis of *Neisseria gonorrhoeae* in the female reproductive tract: neutrophilic host response, sustained infection, and clinical sequelae. Curr Opin Hematol 25: 13–21

Stohl EA, Criss AK, Seifert HS (2005) The transcriptome response of *Neisseria gonorrhoeae* to hydrogen peroxide reveals genes with previously uncharacterized roles in oxidative damage protection. Mol Microbiol 58: 520–532

Stohl EA, Seifert HS (2006) Neisseria gonorrhoeae DNA recombination and repair enzymes protect against oxidative damage caused by hydrogen peroxide. J Bacteriol 188: 7645–7651

Tseng H-J, Srikhanta Y, McEwan AG, Jennings MP (2001) Accumulation of manganese in *Neisseria gonorrhoeae* correlates with resistance to oxidative killing by superoxide anion and is independent of superoxide dismutase activity. Mol Microbiol 40: 1175–1186

Walker E, van Niekerk S, Hanning K, Kelton W, Hicks J (2023) Mechanisms of host manipulation by *Neisseria gonorrhoeae*. Front Microbiol 14: 1119834

Wang Y, Yue X-j, Yuan S-f, Hong Y, Hu W-f, Li Y-z (2021) Internal Promoters and Their Effects on the Transcription of Operon Genes for Epothilone Production in *Myxococcus xanthus*. Front Bioeng Biotechnol Volume 9–2021

Witter AR, Okunnu BM, Berg RE (2016) The Essential Role of Neutrophils during Infection with the Intracellular Bacterial Pathogen Listeria monocytogenes. J Immunol 197: 1557–1565

Zhang HF, Klein Geltink RI, Parker SJ, Sorensen PH (2022) Transsulfuration, minor player or crucial for cysteine homeostasis in cancer. Trends Cell Biol 32: 800–814

